# Identification of *Toxoplasma* Calcium Dependent Protein Kinase 3 as a Stress-Activated Elongation Factor 2 Kinase

**DOI:** 10.1101/2023.03.27.534489

**Authors:** Agnieszka Lis, Carlos Gustavo Baptista, Kelsey Dahlgren, Maria M. Corvi, Ira J. Blader

## Abstract

*Toxoplasma gondii* is an obligate intracellular parasite whose tachyzoite form causes disease via a lytic growth cycle. Its metabolic and cellular pathways are primarily designed to ensure parasite survival within a host cell. But during its lytic cycle tachyzoites are exposed to the extracellular milieu and prolonged exposure requires activation of stress response pathways that include reprogramming the parasite proteome. Regulation of protein synthesis is therefore important for extracellular survival. We previously reported that in extracellularly stressed parasites that the elongation phase of protein synthesis is negatively regulated by the *Toxoplasma* oxygen sensing protein, PhyB. PhyB acts by promoting the activity of elongation factor eEF2, which is a GTPase that catalyzes the transfer of the peptidyl-tRNA from the A site to the P site of the ribosome. In the absence of PhyB, eEF2 is hyper-phosphorylated, which inhibits eEF2 from interacting with the ribosome. eEF2 kinases are atypical calcium-dependent kinases and BLAST analyses revealed the parasite kinase, CDPK3, as the most highly homologous to the *Saccharomyces cerevisiae* eEF2 kinase, *RCK2*. In parasites exposed to extracellular stress, loss of CDPK3 leads to decreased eEF2 phosphorylation and enhanced rates of elongation. Furthermore, co-immunoprecipitation studies revealed that CDPK3 and eEF2 interact in stressed parasites. Since CDPK3 and eEF2 normally localize to the plasma membrane and cytosol, respectively, we investigated how the two can interact. We report that under stress conditions that CDPK3 is not N-myristoylated likely leading to its cytoplasmic localization. In summary, we have identified a novel function for CDPK3 as the first protozoan extracellular-stress induced eEF2 kinase.

**IMPORTANCE:** Here, we identify the first protozoan kinase that phosphorylate elongation factor 2 and demonstrate that it is part of an integrated stress response.

## INTRODUCTION

*Toxoplasma* gondii, which is the causative agent of toxoplasmosis, is an obligate intracellular protozoan parasite that can infect almost all nucleated cells of warm-blooded *mammals* [1]. Upon invasion, the parasite establishes and replicates within a parasitophorous vacuole until it egresses from its host cell and goes on to invade its next host cell [2]. Its success as a pathogen is due to the elaborate mechanisms it uses to complete each step of its lytic cycle as well evade host cell defenses. These include expression of nutrient scavenging and biosynthetic pathways to satisfy its metabolic needs, the deployment of parasite-encoded host cell effectors that alter host signaling, defenses, and transcription, release of receptor complexes into host cell plasma membrane that facilitate host cell invasion, and the ability to develop into a chronic stage infection that is impervious to host immune responses and anti-parasitic agents [3-5].

Like most other obligate intracellular pathogens, host cell egress exposes the parasite to the extracellular milieu. During extended periods of time of extracellular residence, parasite survival requires activation of extracellular stress responses that include alterations in protein abundance [6]. Thus, protein synthesis has emerged as a key extracellular stress responses pathway. Protein synthesis is composed of three stages – initiation, elongation, and termination. Initiation involves ribosome scanning identifying a start methionine ATG codon and the subsequent recruitment of a methionine-charged initiator tRNA to the A site of the ribosome. The initiation factor, eIF-2α, facilitates this process by promoting initiator tRNA association with the ribosome and eIF-2α phosphorylation inhibits its activity. Earlier work revealed that protein synthesis is reduced in parasites exposed to extracellular stress and that this is due, in part, to eIF-2α phosphorylation and inhibition of translation initiation [6].

The *Toxoplasma* oxygen sensing protein, PHYb, is important for efficient parasite lytic growth but dispensable for invasion, replication, or egress. Instead, it is required for extracellular parasite survival and does so promoting translational elongation, which is the process of transferring the peptidyl-tRNA complex from the A site of the ribosome to the P site [7]. The elongation factor, eEF2, catalyzes this process and, like eIF-2α, eEF2 phosphorylation limits its activity [8, 9]. PHYb promotes elongation by limiting eEF2 phosphorylation but how PHYb regulates eEF2 phosphorylation is unknown as is the identify of an eEF2 kinase in *Toxoplasma*. eEF2 kinases are calcium/calmodulin-dependent kinases that have been identified in metazoans and fungi but not protozoans [10]. But their identification is important due to the discovery of eEF2 as a druggable target in *Plasmodium* infections and likely other parasites [11-13].

In *Toxoplasma* and other apicomplexans, CDPKs are the best characterized family of calcium/calmodulin dependent, kinases [14, 15]. The parasite genome predicts the presence of 15 CDPKs with most work focusing on CDPK1 and CDPK3. Both stimulate calcium-dependent exocytosis of the micronemal organelles and egress but only CDPK1 is important for invasion [16-21]. CDPK1 and CDPK3 are both N-myristoylated but only CDPK3 is also palmitoylated and these differences in fatty acylation may be the basis for their differential localization and substrates [19, 22]. But membrane association is not the only feature that defines CDPK3 substrate interactions since putative non-membrane CDPK3 substrates have also been identified [23]. In addition, CDPK3 loss of function mutants inhibits death of extracellular parasites exposed to calcium-ionophore, through an as yet undefined mechanisms [21]. In this study, we identify CDPK3 as an extracellular stress-activated eEF2 kinase that is able to interact with eEF2 in the cytoplasm due to loss of its N-myristoylation. CDPK3 phosphorylation of eEF2 leads to decreases in protein synthesis due to decreased rates of translation elongation suggesting a mechanism by which CDPK3 mediates ionophore-induced extracellular death.

## RESULTS

### eEF2 Phosphorylation During Extracellular Stress is CDPK3 Dependent

Using parasites engineered to conditionally express PHYb via Shield-1 mediated protein stabilization, we reported that decreased PHYb protein levels in parasites exposed to extracellular stress leads to reduced rates of elongation during protein synthesis, and this is a result of increased eEF2 phosphorylation [7]. We therefore hypothesized that PHYb negatively regulates an eEF2 kinase during extracellular stress. eEF2 kinases are calcium-dependent protein kinases that have not yet been identified in protozoans. To first test whether stress-induced eEF2 phosphorylation is Ca^2+^-dependent, PHYb replete and depleted tachyzoites were subjected to extracellular stress for 8 h in the absence or presence of the Ca^2+^ chelator, BAPTA-AM for the final 2 h. Lysates were prepared and Western blotted to detect either total or phosphorylated eEF2. As previously reported, extracellular stress increased eEF2 phosphorylation in the absence of PHYb. But in the presence of BAPTA-AM, stress-induced eEF2 phosphorylation levels were significantly reduced (Fig 1A,B).

**Figure 1:**
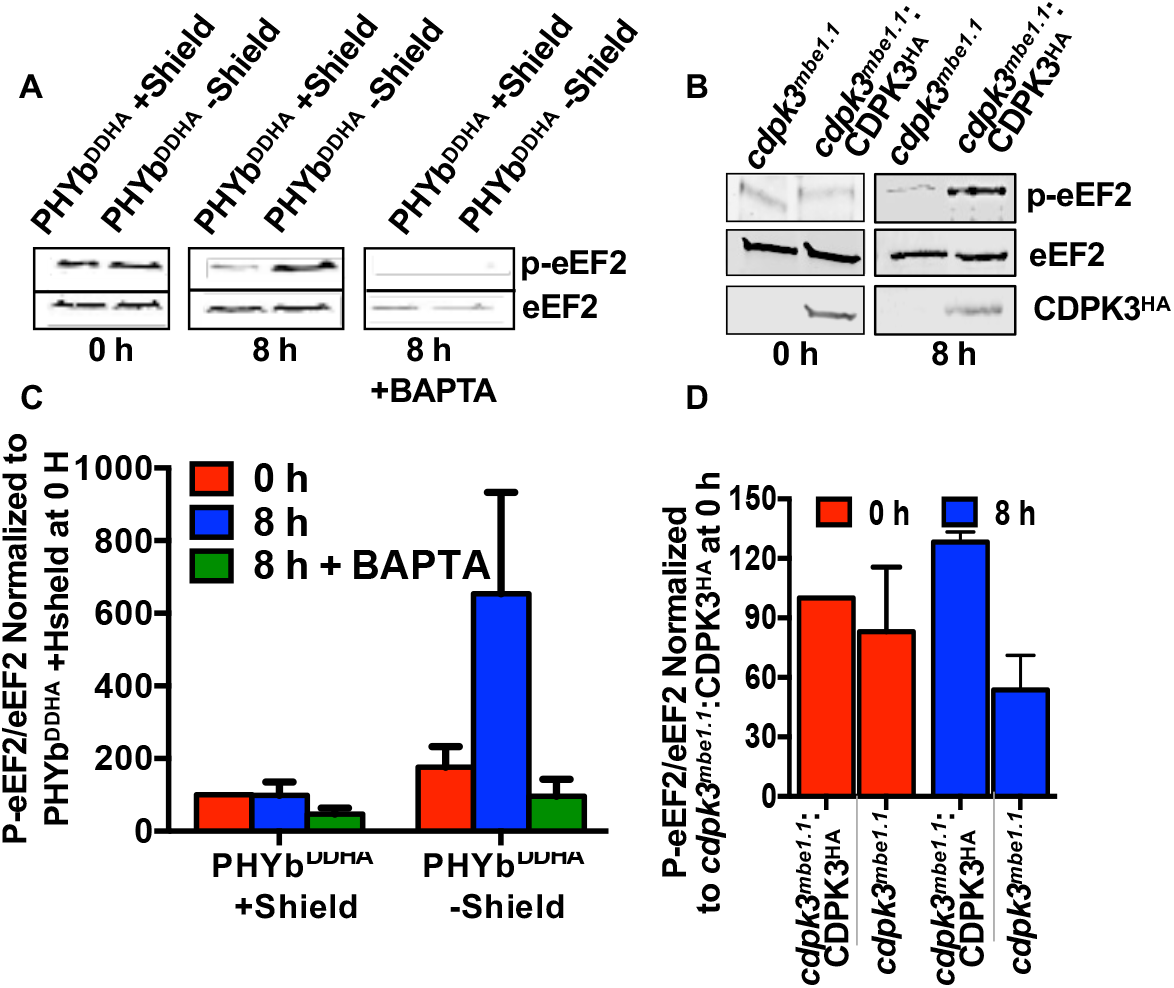
CDPK3 Regulates eEF2 Phosphorylation. **A)** PHYb replete (+Shield) or depleted (–Shield) parasites were either unstressed or extracellularly stressed for 8 h, ±BAPTA-AM for the final 2 h. Lysates were Western blotted to detect total or phospho-eEF2. Shown are representative blots and averages ±SEM of 3 independent experiments. **(B)** CDPK3Δ or CDPK3-overexpressing (CDPK3Δ: CDPK3^HA^) parasites were unstressed or stressed for 8 h. Lysates were blotted to detect total or phospho-eEF2 and HA-tagged CDPK3. Shown are representative blots and the averages ± SD of 3 independent experiments.

eEF2 kinases have been identified in metazoans and fungi but not protozoans. BLAST analysis of human or mouse eEF2 kinases (GenBank Accession numbers AAB58270.1 and AAH60707.1, respectively) against the *Toxoplasma* genome database did not identify any significant hits. The top 5 hits from a BLAST search of the *Saccharomyces cerevisiae* eEF2 kinase, Rck2p [24, 25], were CDPK family members with the regions of homology lying within the conserved protein kinase domain of Rck2p (Table S1 and Fig. S1). Among these, CDPK3 (TgME49_305860) was of interest because it had the highest homology (*E*=2e-^39^) and because the *Plasmodium* homolog of CDPK3 (PfCDPK1) was shown to regulate translation during zygote development in the mosquito midgut [26]. Moreover, *Toxoplasma* mutants harboring loss of function mutations in CDPK3 were resistant to Ca^2+^ ionophore-induced death of extracellular parasites, which was hypothesized to be due to an inability for extracellular parasites to replenish invasion-associated factors that were released following ionophore treatment [20, 21].

We next compared stressed-induced eEF2 phosphorylation levels in *^cdpk3mbe1.1^*, which is parasite strain harboring a point mutation in CDPK3 that inhibits its kinase activity [20, 21]. In these parasites, eEF2 phosphorylation was not increased after exposing the parasites to extracellular stress. In contrast, eEF2 phosphorylation was significantly increased in *cdpk3^mbe1.1^*:CDPK3^HA^, a strain in which wild-type CDPK3 is overexpressed in the mutant strain (Fig 1B,D).

### CDPK3 Regulates Elongation During Protein Synthesis

Extracellular tachyzoites spontaneously secrete host cell adhesins and other factors that function in host cell invasion and other processes and these factors must be replenished for the parasites to remain invasion competent. We previously reported that PHYb-depleted parasites are susceptible to extracellular stress because increased eEF2 phosphorylation decreases rates of elongation during protein synthesis [7]. Elongation rates can be determined by quantifying incorporation of puromycin into nascent polypeptides, which can be detected using puromycin antibody [27]. To assess the impact CDPK3 on elongation, *cdpk3^mbe1.1^*and *cdpk3^mbe1.1^*:CDPK3^HA^ parasites were incubated extracellularly for 4 or 8 hours and then puromycin was added for increasing amounts of time. Lysates were prepared, separated by SDS-PAGE, and Western blotted to detect puromycylated-peptides. We observed that extracellular stress increased rates of elongation only in *cdpk3^mbe1.1^* but not *cdpk3^mbe1.1^*:CDPK3^HA^ parasites (Fig. 2). It is important to note that elongation was not as severely inhibited in the *cdpk3^mbe1.1^*:CDPK3^HA^ strain as we previously reported for the parental RH strain parasites, which is likely due to CDPK3 overexpression in these parasites.

**Figure 2:**
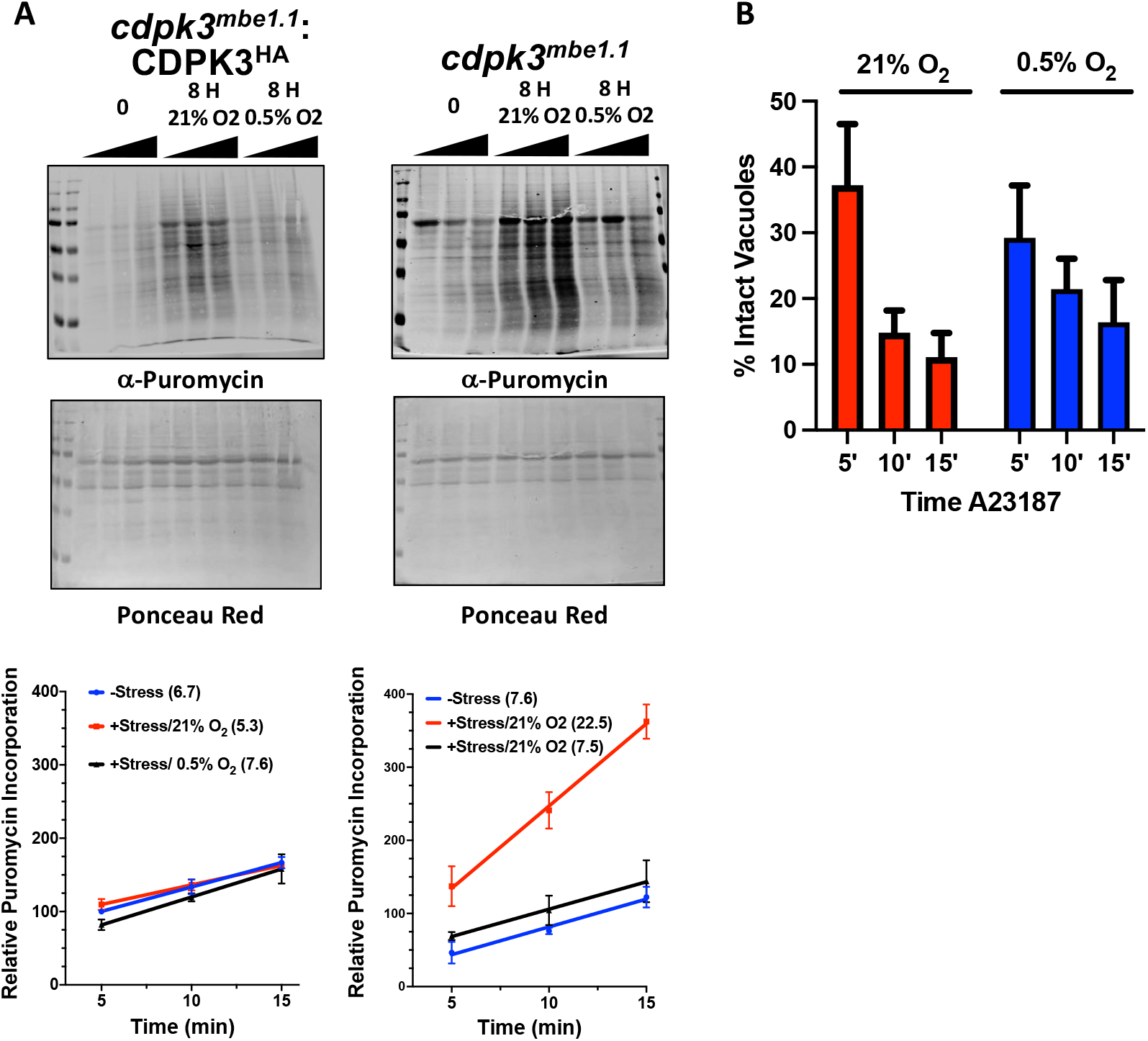
CDPK3 Negatively Regulates Translation Elongation. **A)** Puromyclated peptides were detected by Western blotting lysates prepared from either freshly egressed parasites or parasites incubated extracellularly for 8 h at 21% O_2_ or 0.5% O_2_ were incubated with puromycin for for the indicated times. Lysates were prepared, separated by SDS-PAGE and then either Western blotted to detect puromyclated peptides (top) or stained with Ponceau (bottom). Shown are representative blots from 3 independent experiments. Graphs below show averages ± standard deviations of elative puromycin/ponceau labelling intensities determined for each time point. The slope of each line was calculated to determine rate of puromycin incorporation and shown in graph legend as number following each condition. **B)** *cdpk3^mbe1.1^*:CDPK3^HA^ parasites were grown for on coverslips for 30 h at 21% or 0.5% O_2_. A23187 was then added for the indicated times and the cells were immediately fixed and stained to detect SAG1 by immunofluorescence. Shown are averages ± standard deviations of 2 experiments performed in triplicate with at least 50 random vacuoles counted/coverslip.

Previously, we reported that translation elongation was inhibited in parasites exposed to extracellular stress at 21% O_2_ but not at 0.5% O_2_ and that this phenotype was mediated by PHYb [7]. Thus, we used puromycin incorporation to assess translation elongation at 0.5% O_2_. At low O_2_, rates of translation elongation were not changed in either CDPK3-deficient or overexpressing parasites (Fig. 2A). Taken together, these data indicate that CDPK3 regulates translation elongation as part of an extracellular stress response that functions upstream of O_2_/PHYb signaling.

### Extracellular Stress Promotes CDPK3-eEF2 Interactions

Having established that extracellular stress increased CDPK3-dependent eEF2 phosphorylation, we next sought to determine whether the two proteins interact. We prepared extracts from either freshly egressed or extracellularly stressed *cdpk3^mbe1.1^* and *cdpk3^mbe1.1^*:CDPK3^HA^ parasites. Lysates were then incubated with anti-HA antibodies, immune complexes separated by SDS-PAGE, and CDPK3^HA^ and eEF2 detected by Western blotting. It was observed that eEF2 only interacted with CDPK3 in stressed parasites (Fig. 3A).

**Figure 3:**
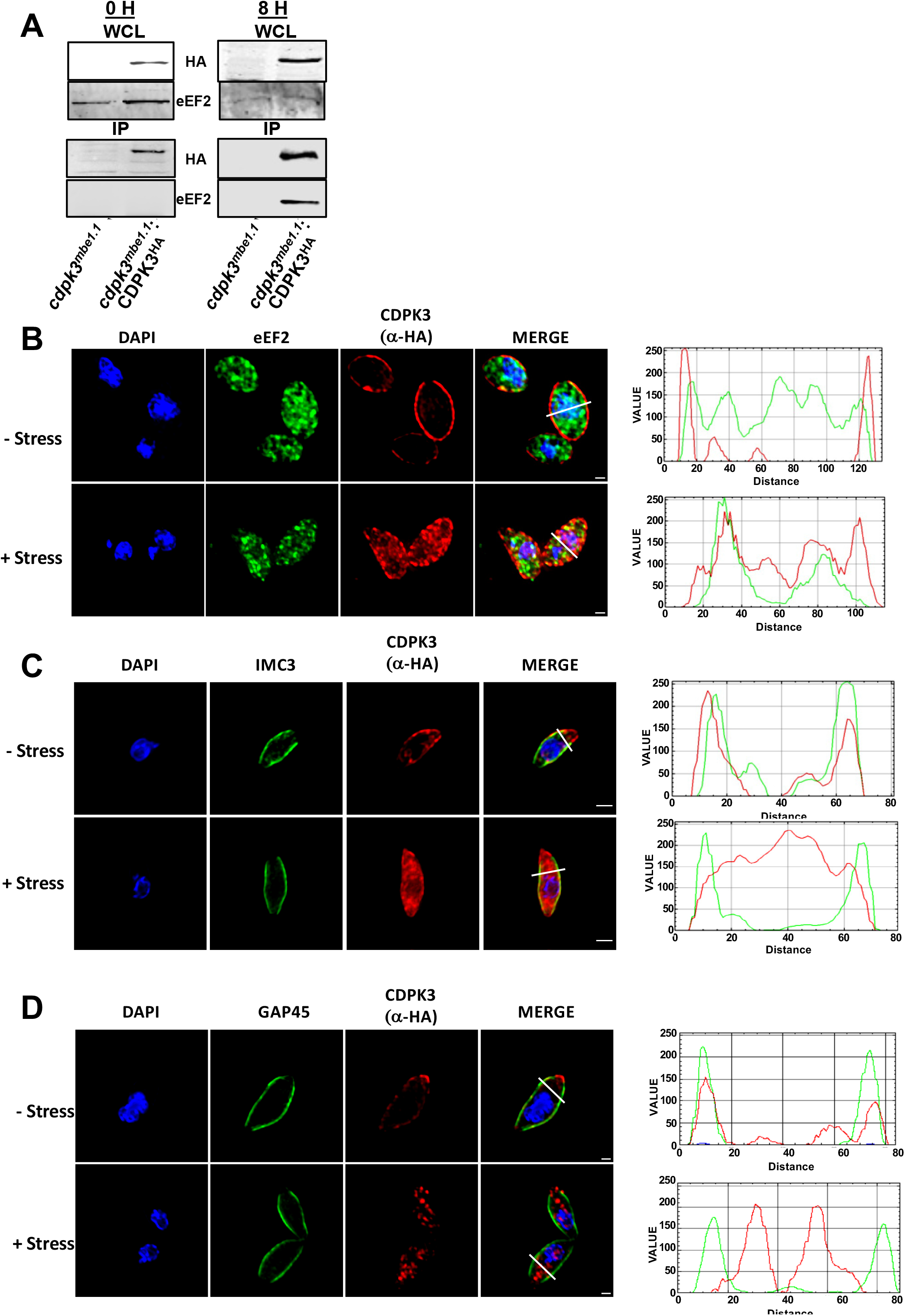
Extracellular Stress Promotes CDPK3/eEF2 Interactions. **A)** CDPK3Δ or CDPK3Δ:CDPK3^HA^ parasites were incubated extracellularly for 0 or 8 h. Whole cell lysates were prepared and either Western blotted to detect CDPK3 and eEF2 or incubated with anti-HA/Protein A beads and the immune complexes blotted to detect CDPK3^HA^ and eEF2. Shown are representative blots from three independent experiments. **B-D)** CDPK3Δ:CDPK3^HA^ parasites were incubated extracellularly for 0 or 8 h and then stained to detect CDPK3^HA^, eEF2, IMC3, or GAP45. Plots on right represent staining intensities of each protein as shown by the white line in the overlay images. Shown are representative images. Scale bar = 1 μm.

In neurons and other cell types, eEF2 is a cytoplasmically-enriched, soluble nucleocytoplasmic protein [28-30] and HYPERLOPIT analyses predicted *Toxoplasma* eEF2 to also be a highly enriched in the cytoplasm [31]. On the other hand, CDPK3 is embedded in the inner leaflet of the parasite’s plasma membrane [19, 21]. Because the inner membrane complex (IMC) is directly apposed to the plasma membrane, cytoplasmic proteins do not have direct access to the plasma membrane. Thus, for CDPK3 and eEF2 to interact, one of the proteins would likely have to alter its localization. We therefore used immunofluorescence assays to examine localization of each protein in freshly egressed and stressed parasites. eEF2 was observed to maintain its cytoplasmic localization regardless of stress (Fig. 3B). On the other hand, extracellular stress induced a significant fraction of CDPK3 to relocalize from the plasma membrane to the cytoplasm. The altered localization was not due to a breakdown of the parasite plasma membrane or underlying IMC as localization of either GAP45 (plasma membrane associated) or IMC3 (a resident IMC protein) were unaltered by stress (Fig. 3 C&D).

### Extracellular Stress Alters CDPK3 N-Myristoylation

CDPK3 is embedded in the plasma membrane by virtue of its myristoylation and palmitoylation at positions ^2^Gly and ^3^Cys, respectively [18, 19, 21, 32]. We therefore hypothesized that stress-induced CDPK3 cytoplasmic localization was due to alterations in its fatty acylation. To test this hypothesis, stressed and unstressed *cdpk3^mbe1.1^* and *cdpk3^mbe1.1^*:CDPK3^HA^ parasites were incubated in the presence of azido-myristic acid or azido-palmitic acid for 2 hours. Lysates were then prepared and incubated with alkyne-conjugated biotin, separated by SDS-PAGE, and blotted to detect total protein fatty acylation. We found that extracellular stress had no significant impact on either global myristoylation or palmitoylation (Fig. 4A&B). Next, CDPK3^HA^ was immunoprecipitated from the azido-labelled lysates and the immune complexes were Western blotted using streptavidin to specifically assess CDPK3 myristoylation or palmitoylation. We found that while extracellular stress had no apparent impact on its palmitoylation, CDPK3 myristoylation was significantly reduced (Fig. 4 C&D).

**Figure 4:**
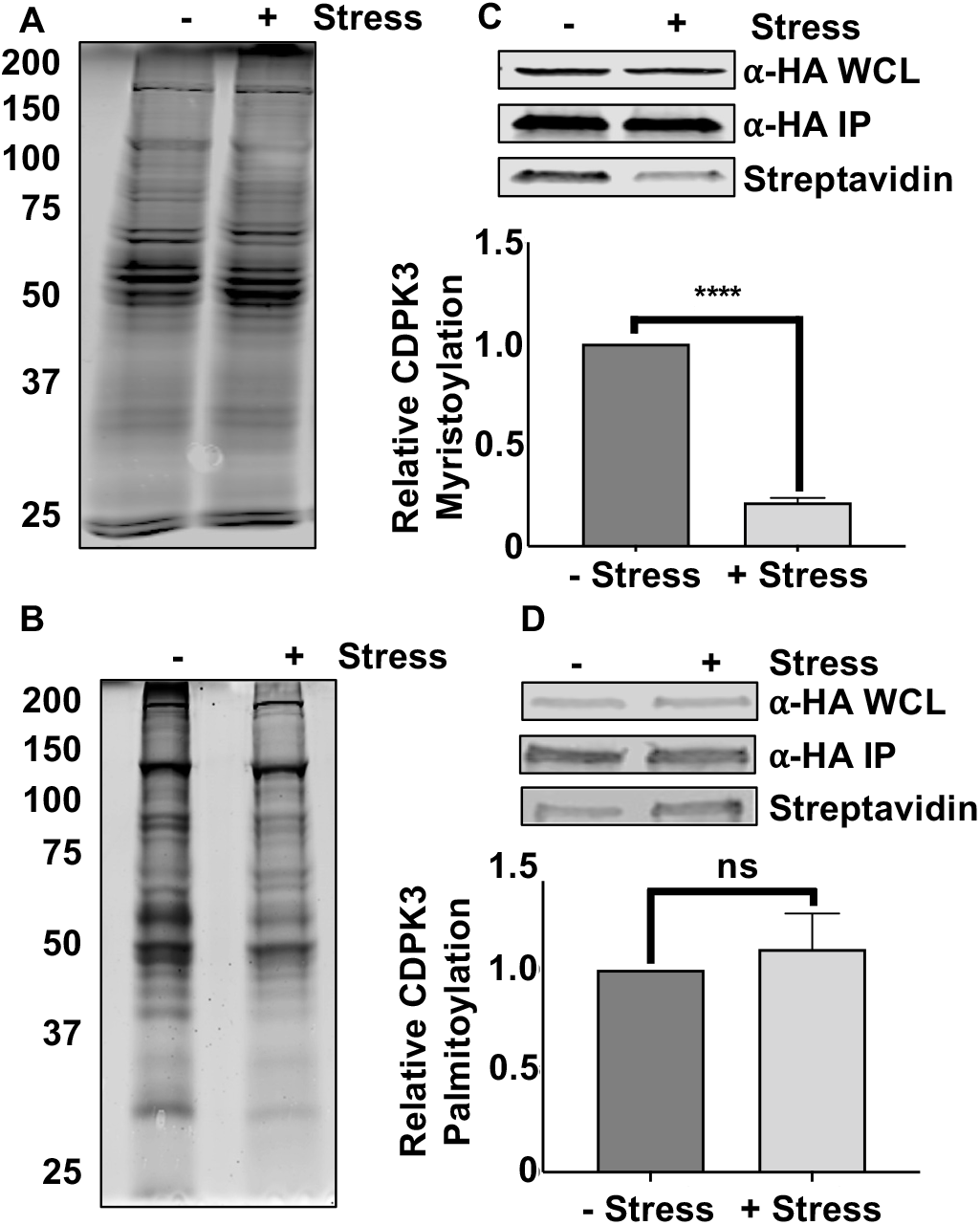
Extracellular Stress Alters CDPK3 Myristoylation. Freshly egressed (-stress) or extracellularly stressed (+stress) CDPK3Δ:CDPK3^HA^ parasites were incubated with azido-myristate **(A,C)** or azido-palmitate **(B,D)** for 1 hour. Whole cell lysates (1/10 or total) were incubated with alkyne-biotin, separated by SDS-PAGE, and blotted with streptavidin to detect total myristoylated **(A)** or palmitoylated **(B)** proteins or total CDPK3 (⍺-HA WCL in **C&D**). The remainder of lysates were incubated with anti-HA beads to capture CDPK3^HA^ and immune complexes incubated with alkyne-biotin to biotinylate the myristate **(C)** or palmitate **(D)** analogs. The reactions were separated by SDS-PAGE and Western blotted to detect total CDPK3 (αHA) or and acylated CDPK3 (streptavidin). Shown are representative blots from three independent experiments and graphs represent average ± SD of the ratios of fatty acylated CDPK3^HA^:total CDPK3^HA^.

N-myristoylation is catalyzed by protein N-myristoyltransferase (NMTase) that in *Toxoplasma* is encoded by a single essential gene (TGME49_209160). TgNMTase is predicted to be important for tachyzoite fitness [33] and migrates as a ∼57 kDa protein in SDS-PAGE [34]. Western blot analysis indicated that decreased CDPK3 myristoylation was not due to a stress-induced decrease in TgNMTase expression, which is consistent with the observation that extracellular stress did not have an apparent global impact on protein myristoylation (Fig. 5A). The NMTase reaction is initiated by the enzyme first complexing with myristoyl-CoA followed by binding of the protein substrate whose ^2^Gly has been exposed by release of the N-terminal methionine by a methionine aminopeptidase. A series of conformational changes brings the reactants near enough to allow conjugation of the myristic acid to ^2^Gly after which the myristoylated protein is released. We therefore assessed whether extracellular stress affected CDPK3/TgNMTase interactions by immunoprecipitating CDPK3^HAHA^ from unstressed and extracellularly-stressed parasites and then Western blotted to detect CDPK3^HA^ or TgNMTase. In unstressed parasites, the enzyme:substrate complex was undetectable indicating that the N-myristoylation reaction was efficient (Fig. 5). In contrast, TgNMTase could be detected in the CDPK3^HA^ immunoprecipitates from the stressed parasites. These data together suggest that defects in CDPK3 myristoylation was a due to TgNMTase being unable to properly interact with CDPK3.

**Figure 5:**
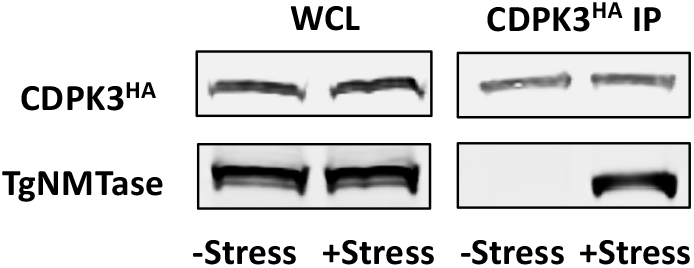
CDPK3/TgNMTase Interactions are Enhanced in Parasites Exposed ot Extracellular Stress. Lysates from freshly egressed (-stress) or extracellularly stressed (+stress) CDPK3Δ:CDPK3^HA^ parasites were either Western blotted to detect total CDPK3^HA^ and TgNMTase or incubated with anti-HA/Protein A beads and the immune complexes blotted to detect CDPK3^HA^ and TgNMTase. Shown are representative blots from three independent experiments.

## DISCUSSION

Metabolic and other essential processes of obligate intracellular pathogens are designed for their intracellular lifestyle. But these pathogens are exposed to the extracellular milieu as they move between host cells and how they survive under these conditions is largely unknown. In a previous study, we identified elongation as a process that is modulated as part of this stress response. We further demonstrated that eEF2 phosphorylation is negatively regulated by PHYb and that this negatively regulatory loop is required to allow parasites to persist under prolonged exposure to extracellular stress. In this study, we sought to identify the stress-activated eEF2 kinase and based upon results from co-immunoprecipitation, eEF2 phosphorylation status, and localization studies we conclude that CDPK3 serves this function.

CDPK3 was first identified in a genetic screen designed to identify parasite mutants that were resistant to parasites egressing from host cells upon exposure to the Ca^2+^ ionophore A23187 [20, 21]. In addition, this mutant (named MBE1.1) were also resistant parasite death induced when extracellular parasites were exposed for prolonged times to the Ca^2+^ ionophore. CDPK3 substrates have been identified including a myosin isoform whose phosphorylation likely underlies the egress phenotype [23, 35]. But neither microneme secretion nor motility were affected in extracellular CDPK3 mutants leading to the suggestion that the CDPK3 mutants are unable to replenish those invasion-associated factors secreted upon Ca^2+^ ionophore exposure [20, 21]. Our findings here support such a model in which CDPK3 negatively regulates protein synthesis in extracellular parasites and that loss of CDPK3 removes this repression.

Stress responses modulate protein synthesis to allow a cell to reprogram its proteome. In most organisms including *Toxoplasma* and other apicomplexans, translation initiation is the best characterized mechanism for stress responses to regulate protein synthesis [36] with eIF2α phosphorylation as a key target. In *Toxoplasma*, distinct kinases phosphorylate eIF2α in response to oxidative, endoplasmic reticulum, nutritional, and extracellular stresses [37-40]. Elongation, on the other hand, is a less well understood though recognized stress prokaryotic and eukaryotic response target [41-43]. Unlike eIF2α as the primary target for regulating translation initiation, numerous elongation factors can be modulated by stress and these include eEF2 [7, 43-47]. Our identification of CDPK3 as an eEF2 kinase represents, to our knowledge, the first protozoan eEF2 kinase. Our data showing that CDPK3 only impacts eEF2 phosphorylation in extracellularly stressed parasites suggests the presence of additional eEF2 kinases. Whether other CDPK family members serve this function remains to be determined but given that eEF2 kinases are not essential in *Saccharomyces*, C. elegans, or mice [48, 49] we predict that like CDPK3 that non-essential *Toxoplasma* CDPKs would most likely act as eEF2 kinases. Identification of these kinases is important given the increasing focus on eEF2 as a drug target in related parasites such as *Plasmodium* and *Eimeria* [12, 13, 49].

The IMC in *Toxoplasma* and other apicomplexans is a specialized double membrane organelle that lies directly underneath the plasma membrane and functions in parasite motility, secretion, and replication [2, 50-52]. Its close apposition to the plasma membrane limits contact between plasma membrane and cytosolic factors. Thus, we predicted that marked relocalization of either eEF2 or CDPK3 would be required for the two proteins to interact. Our finding CDPK3 that relocalizes under extracellular stress was reminiscent of glycolytic enzyme relocalization in extracellular parasites [53], although there are significant differences exist between the two. First, glycolytic enzymes relocalize from the cytosol to the periphery. Second, glycolytic enzymes associate with membranes via electrostatic interactions rather that embedment via fatty acylation. Finally, glycolytic enzymes associate with the cytoplasmic face of the IMC, a finding further supporting our observations that the IMC remains intact upon exposure to extracellular stress.

Mechanistically, CDPK3 relocalization is likely the result of newly synthesized CDPK3 from becoming N-myristoylated since N-myristoylation is considered an irreversible posttranslational modification[54]. On the other hand, we found that CDPK3 was palmitoylated, which was initially surprising given that S-palmityl acyltransferase (PAT) are membrane associated and ^2^Gly myristoylation is thought to mediate PAT access to N-terminal cysteines in substrates, like CDPK3 that is palmitoylated at ^3^Cys [19, 21]. One explanation for this finding is that ^3^Cys-palmitoylation is independent of ^2^Gly myristoylation. Alternatively, CDPK3 was reported to also be Ser-palmitoylated [32] and our assays cannot discriminate between single of multiple palmitoylation events nor can it discriminate between the modified amino acids. Regardless, these data suggest that CDPK3 utilizes multiple membrane targeting mechanisms that is likely a binding protein since the protein lacks apparent regions of positively charged amino acids.

The ability for CDPK3 and NMTase to interact in response to extracellular stress suggests that CDPK3 becomes “trapped” with NMTase in an intermediate complex. Based on structural studies of the NMTase reaction [55], we hypothesize that CDPK3 associates with NMTase-bound myristyl-CoA in a manner that ^2^Gly cannot interact with the carbonyl group of myristyl-CoA. It is also unclear what triggers this block in CDPK3 N-myristoylation. Our future studies will address these questions.

Translation elongation is a recognized target for hypoxic stress responses [56, 57] and in *Toxoplasma* our collective data points to a model in which PHYb negatively regulates CDPK3 phosphorylation of eEF2. This is most likely achieved by CDPK3 and/or eEF2 serving as substrates for PHYb’s prolyl hydroxylase activity. In murine cells, eEF2 kinase is a prolyl hydroxylase substrate and hydroxylation of ^98^Pro blocks eEF2 kinase activity [58]. This would contrast with *Toxoplasma* PHYb that is active at high oxygen and thus PHYb-mediated prolyl hydroxylation of CDPK3 would be expected to promote CDPK3 activity. It is also possible that PHYb acts by preventing CDPK3 from binding its substrates or by promoting the activity of an eEF2 phosphatase.

In summary, we identified CDPK3 as a kinase that regulates the phosphorylation state and activity of eEF2 in response to *Toxoplasma* exposure to extracellular stress. We further determined that interactions between CDPK3 and eEF2 are mediated by reduced N-myristoylation and that is due an apparent defect in the NMTase reaction. Given the conservation of CDPK3 and eEF2 amongst other apicomplexan parasites, we expect that this signaling axis will be conserved in these parasites as well.

## MATERIALS AND METHODS

### Cells and Parasites

The type I *ckpk3^mbe1.1^* and *ckpk3^mbe1.1^:CDPK3^HA^ Toxoplasma* strains [21] were obtained from Dr. Gustavo Arrizabalaga (Indiana University School of Medicine) and maintained by passage in human foreskin fibroblasts (HFFs) in complete medium (Dulbecco’s Modified Eagle Medium supplemented with 10% heat-inactivated fetal calf serum, L-glutamine, and penicillin/streptomycin) as previously described [59]. Parasites and host cells were regularly tested for Mycoplasma contamination using the Mycoalert detection kit (Lonza; Basel, Switzerland). For all assays, parasites harvested from unlysed HFF monolayers were released by syringe lysis by passage through a 27-gauge needle followed by washing in serum-free media. Extracellular stress was applied by incubating the parasites in complete medium at 37°C at either 21% O_2_ or 0.5% O_2_ (Invivo2 Baker, Sanford, ME). Calcium-ionophore induced egress assays were performed essentially as described [20]. Briefly, parasites were grown in HFFs on coverslips for 30 hours at 21% or 0.5% O_2_. The medium was then replaced with prewarmed Hanks buffered saline solution containing 8 μM A23187 for the indicated times and then ice cold methanol was added to fix the cells. Lysed and intact vacuoles were identified by immunofluorescence staining of the parasite surface protein SAG1 (Novus; Centennial, CO).

### Western Blotting

Parasites were pelleted by centrifugation 2000 *g* for 8 min at 4°C and lysed in boiling SDS-PAGE sample buffer containing 2-β mercaptoethanol. Equivalent cell volume of lysates were separated by SDS-PAGE gels, transferred to nitrocellulose membranes and blocked with Odyssey Blocking Buffer (LI-COR Biosciences, Lincoln, NE), blotted using antibodies to detect total eEF2 (Abcam #33523; Cambridge, UK) phospho-eEF2 (Cell Signaling #2331S; Danvers, MA), anti-HA (Sigma #12CA5; St. Louis, MO), and TgNMTase [34]. Blots were imaged using a LI-COR Odyssey scanner and analyzed using Image Studio software (LI-COR; Omaha, NE).

### Immunoprecipitation

Freshly egressed or extracellularly stressed parasites were lysed in 50 mM Tris-HCl pH 7.4 with 1% Triton X-100, 150 mM NaCl, 1 mM NaF, 1 mM CaCl_2_, 0.2 mM Na3VO4, 1X protease inhibitor cocktail (Thermo Fisher; Waltham, MA), incubated on ice for 30 min and then subjected to sonication. Lysates were clarified by centrifugation at 16,000 *x*g, incubated with mouse α-anti-HA clone 12CA5 conjugated with agarose beads (Abcam #214758) for 16 hours at 4°C. Immune complexes were separated by SDS-PAGE, transferred to nitrocellulose, and processed for Western blotting.

### Immunofluorescence Microscopy

Freshly egressed or extracellularly stressed parasites (10^7^) were fixed with 4% w/v paraformaldehyde in phosphate-buffered saline (PBS) for 20’ at room temperature, permeabilized with 0.1% Triton X-100 in PBS for 10’, and blocked 5% w/v bovine serum albumin (BSA) in PBS for 20’. Parasites were then incubated with a combination of antibodies to detect HA, and either eEF2, IMC3 (from Dr. Marc-Jan Gubbels; Boston College, MA) or GAP45 (gift from Dr Gary Ward; University of Vermont, VT) for 16 hours at 4°C and followed by secondary Alexa-Fluor 488- or Alexa Fluor-594-conjugated antibodies for another 60 min (Thermo Fisher Scientific).

For image acquisition, a 0.2-μm z-increment serial image stacks were collected with capture times from 100 to 500 msec (100X Plan Apo oil immersion lens, 1.46 numerical aperture) on a motorized Zeiss Axioimager M2 stand equipped with a rear-mounted excitation filter wheel, a triple pass (DAPI/fluorescein isothiocyanate [FITC]/Texas Red) emission cube, differential interference contrast (DIC) optics, and an Orca ER charge-coupled-device (CCD) camera (Hamamatsu, Bridgewater, NJ). Images were collected with Volocity, version 6.1, Acquisition Module (Improvision Inc., Lexington, MA). Individual channel stacks were deconvolved by a constrained iterative algorithm, pseudo colored, and merged using the Volocity, version 6.1, Restoration Module. All images presented here are summed stack projections of merged channels.

### Translation Elongation Assay

Puromycin incorporation was performed essentially as described [7]. Briefly, freshly harvested parasites (3×10^6^) were resuspended in 1 ml of complete DMEM and incubated at 21% or 0.5% O_2_ for 0 or 8 h at 37°C. Puromycin (10 μg/ml) (Sigma) was added for the indicated times and the parasites were pelleted, washed in ice-cold PBS, and analyzed by Western blotting using an antipuromycin antibody. Each lane was scanned and quantified densitometrically. Puromycin incorporation was normalized to the Ponceau S stain intensity of the membranes, and translation rates were calculated from slopes of the lines generated using linear regression analysis with Prism Software (GraphPad, La Jolla, CA).

### Fatty Acylation Assays

CDPK3 myristoylation and palmitoylation was assessed using the CLICK-iT kit following the manufacturer’s protocol (ThermoFisher). Briefly, intracellular (-stress) or freshly egressed (+stress) parasites were incubated with Click-iT™ myristic acid azide (25 µM) or palmitic acid azide (50 µM) for 4 hours at 37°C. Parasites were collected and lysed by sonication in RIPA buffer (150 mM NaCl, 50 mM Tris-HCl pH 7.4, 10 mM MgCl, 0.5% sodium deoxycholate, 0.1% SDS, and 1% Triton with protease inhibitor cocktail (Roche)). To assess global fatty acylation, lysates were incubated with biotin alkyne (20 µM) using in the Click-iT™ Protein Reaction Buffer protocol and the proteins were precipitated and then resuspended and boiled for 10 minutes in 2X Laemmli buffer. To assess CDPK3 fatty acylation, lysates were incubated with rabbit α-HA antibody and then caputured using Protein G Agarose (Pierce; Rockford, IL). Immunecomplexes were then incubated with biotin alkyne (20 µM) in the Click-iT™ Protein Reaction Buffer Kit. Lysates and immunecomplexes were separated by SDS-PAGE and blotted to detect either biotynlated proteins (IRDye® 680LT streptavidin (Li-Cor)) or CDPK3^HA^ (rat-anti-HA)/anti-rat DyLight™ 800 (Invitrogen).

### Statistics

Unless otherwise noted, all experiments were repeated a minimum of three times. Data were expressed as mean ± SD, plotted and tested for significance using two-tailed Student’s *t*-test with Prism Software (GraphPad, La Jolla, CA).

## ACKNOWLEDGEMENTS

We thank Drs. Gustavo Arrizabalaga, Gary Ward, and Marc Jan Gubbels for supplying critical reagents and members of our lab for helpful discussions

## FIGURE LEGENDS

**Figure S1: Alignment of Top 24 Hits from Yeast RCK2 BLAST Analysis Against *Toxoplasma* Genome.**

**Table S1: Results of Yeast RCK2 BLAST Analysis Against Toxoplasma Genome.**

